# Filamentous cheater phages drive bacterial and phage populations to lower fitness

**DOI:** 10.1101/2025.04.01.646652

**Authors:** Nanami Kubota, Michelle R. Scribner, Vaughn S. Cooper

**Affiliations:** Department of Microbiology and Molecular Genetics, School of Medicine, University of Pittsburgh, Pittsburgh, Pennsylvania 15219, USA; Center for Evolutionary Biology and Medicine, University of Pittsburgh, Pittsburgh, Pennsylvania 15219, USA

**Keywords:** *Pseudomonas aeruginosa*, filamentous phage, Inoviridae, cheating, Tragedy of the Commons, experimental evolution, lysogeny, Pf phage, intracellular competition

## Abstract

Many bacteria carry phage genome(s) in their chromosome, which intertwines the fitness of the bacterium and the phage. Most *Pseudomonas aeruginosa* strains carry filamentous phages called Pf that establish chronic infections and do not require host lysis to spread. However, spontaneous mutations in the Pf repressor gene (*pf5r*) can allow extreme phage production that slows bacterial growth and increases cell death, violating an apparent détente between bacterium and phage. We observed this paradoxical outcome in an evolution experiment with *P. aeruginosa* in media simulating nutrients from the cystic fibrosis airway. Bacteria containing *pf5r* mutant phage grow to a lower density but directly outcompete their ancestor and convert them into *pf5r* mutants via phage superinfection. Reduced fitness therefore spreads throughout the bacterial population, driven by weaponized Pf. Yet high intracellular phage replication facilitates another evolutionary conflict: “cheater miniphages” lacking capsid genes and the superinfection exclusion gene (*pfsE*) invade populations of full-length phages within cells. Although bacteria containing both full-length phages and miniphages are most immune to superinfection by limiting the Pf receptor, this hybrid vigor is extremely unstable, as a classic Tragedy of the Commons scenario ensues that causes complete prophage loss. The entire cycle – from phage hyperactivation to miniphage invasion to prophage loss – can occur within 24h, showcasing rapid coevolution between bacteria and their filamentous phages. This study demonstrates that *P. aeruginosa*, and potentially many other bacterial species that carry filamentous prophages, risk being exploited by these phages in a runaway process that reduces fitness of both host and virus.

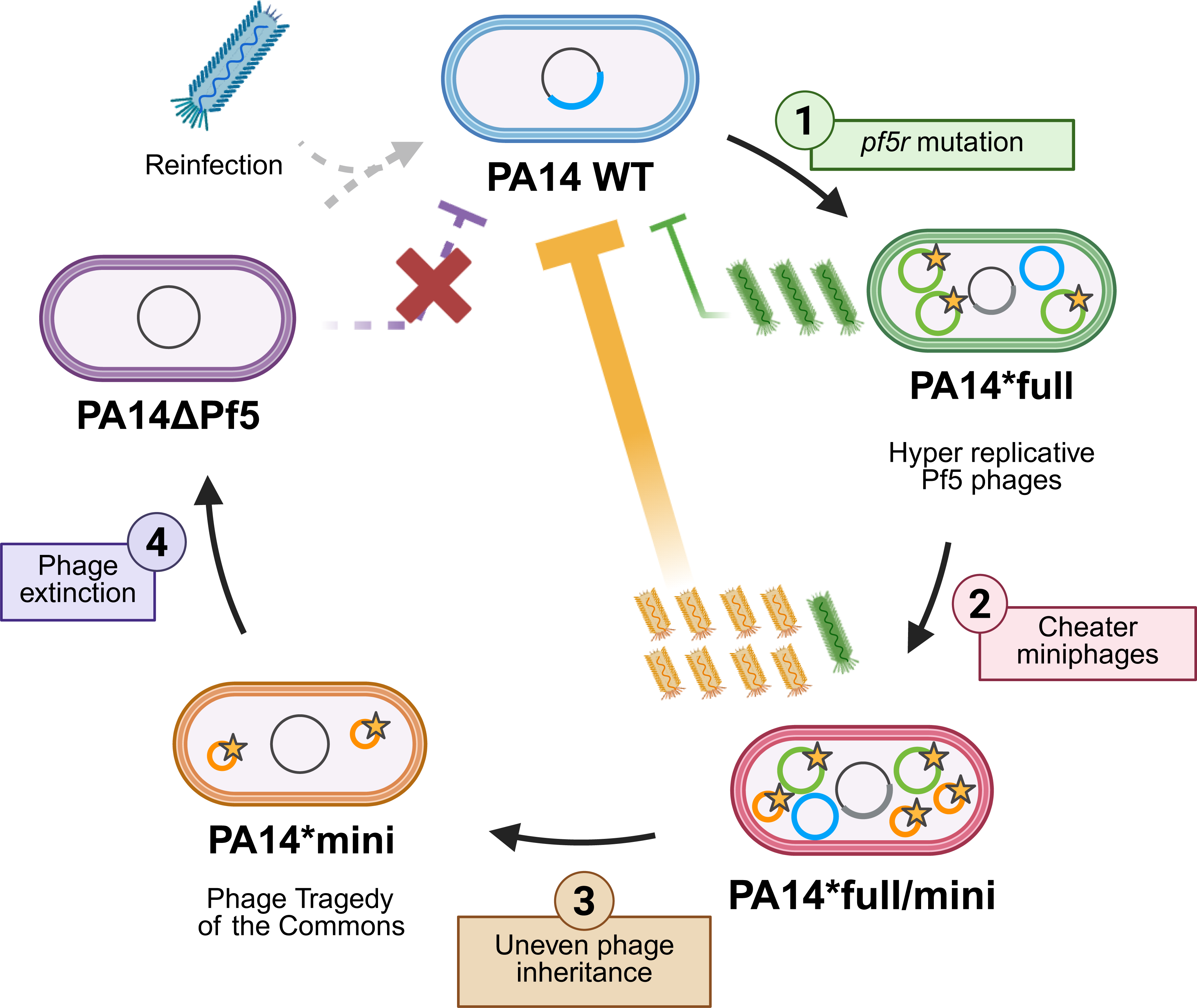

## Introduction

Many bacteria carry genetic parasites such as lysogenized phages that inevitably influence the evolutionary trajectory of their bacterial host as well as the phage. The fitness of temperate phages that are integrated into their bacterial host chromosome (prophages) is tied to their host, and likewise, the host phenotype and fitness are also influenced by the presence of prophage.^1–4^ While prophages may encode beneficial traits for the bacteria, such as antibiotic resistance genes^5,6^ or superinfection exclusion mechanisms^7^, maintenance of an active prophage encumbers costs such as the potential for cell lysis.^6^ Filamentous phages belonging to the *Inoviridae* family are unique from other phages because they do not require host cell lysis to produce progeny.^8,9^ This lowers the risk to the host to maintain an active prophage. Since phage activation and replication do not necessarily kill the host, filamentous phage can achieve increased population size, experience more mutations, and develop greater diversity. Consequently, seemingly clonal bacterial cells may differ in the genetic and phenotypic composition of their phage populations, broadening heterogeneity and the number of traits that selection may act upon.

Many *Pseudomonas aeruginosa* strains carry filamentous Pf prophage or Pf phage-like elements in their chromosome^10^, including common laboratory strains like PAO1 with its Pf4 and Pf6 prophages and UCBPP-PA14 (shortened as PA14) with its Pf5 prophage.^11^ *P. aeruginosa* is a Gram-negative opportunistic pathogen that causes many hospital-acquired infections and is the major infectious threat for persons with cystic fibrosis (CF). As of 2022, approximately 26.0% of people in the CF Foundation Patient Registry tested positive for *P. aeruginosa* with that percentage increasing to 59.5% in those who met the criteria for Advanced Lung Disease.^12^ As CF patients age, their infecting *P. aeruginosa* isolates are more likely to carry Pf prophage.^10^ This association may be driven by functions provided by Pf phages to their bacterial host during infection, as their negatively charged filaments and ability to form liquid crystalline structures can make biofilms more robust, delay wound healing, and protect bacteria from antibiotics.^3,10,13–15^ Consequently there are both fundamental and clinical needs to understand ecological and evolutionary interactions between Pf and their host bacteria.

In a previous evolution experiment with *P. aeruginosa* PA14 grown in media containing nutrients found in CF lung environments, we observed that this nutritional environment was sufficient to select for clinically relevant bacterial phenotypes.^16^ Here we identify that within one of the evolved populations, mutations in the Pf5 phage repressor (*pf5r*) gene within the Pf5 prophage genome rose to high frequency. The Pf phage repressor functions by repressing the transcription of Pf phage excise element (*xisF5*), preventing Pf prophage from leaving the bacterial chromosome.^17^ Previous studies with a better-studied Pf phage, Pf4, showed that mutations in the Pf4 phage repressor (*pf4r*) gene or promoter region can lead to emergence of superinfective, hyper-replicative Pf4.^18–22^ Furthermore, overexpression of the Pf4 excise element (*xisF4*) is correlated with phage virion production.^19^ Here we find that the *pf5r* mutation converts Pf5 phage into a “superinfective” form, capable of infecting wildtype (WT) PA14 already lysogenized by WT Pf5 prophage and increasing Pf5 phage replication and production. However, infected cells grew significantly slower than those with WT Pf5, posing a puzzle of how the *pf5r* mutation rose to high frequency.

We show that PA14 *pf5r* mutants outcompete their ancestor by weaponizing their *pf5r* mutant phage and demonstrate that this imposes a greater cost on the newly superinfected victim cell than on the *pf5r* phage donor cell. Paradoxically, this dynamic progressively reduces bacterial population fitness. Yet these selfish superinfective phage themselves become victimized by cheater “miniphages” lacking structural genes and the Pf superinfection exclusion (*pfsE*) gene, rapidly producing a phage Tragedy of the Commons that leads to Pf5 loss as well as a host susceptible to reinfection. The robust conditions of this experimental model indicate the potential for rapid coevolution between bacteria and filamentous phage in *P. aeruginosa,* as well as in other bacteria carrying inoviruses, affecting a range of traits essential for fitness.

## Results

### Defective Pf phage repressor leads to runaway Pf replication and phage production

Mutations that disrupt the prophage repressor gene can trigger phage activation, leading to the emergence of hyper-replicative Pf phage.^17,20,22^ In a previous evolution experiment with *P. aeruginosa* PA14 that was analyzed by longitudinal whole-population genome sequencing^16,23^, we noticed increased sequencing coverage, or read depth, along the Pf5 prophage region in one evolved population (Figure S1). Further investigation of sequencing results showed a 16 bp duplication mutation within the Pf5 phage repressor gene (*pf5r*) and circularization of the Pf5 genome, indicating active Pf5 phage (Table S1). The *pf5r* gene is within the prophage region that is integrated into the bacterial chromosome, and so we denote bacteria and phage genotypes with the *pf5r* mutation using an asterisk (e.g., PA14* and Pf5*, respectively, see Methods and Table S2 for additional nomenclature). To isolate the *pf5r* mutation in a clean genomic background, we attempted to infect PA14 wildtype (WT) with the Pf5* phage from the evolution experiment and were successful (see Methods for more detail). The prophage genome and flanking bacterial genes are shown in Figure 1A. Compared to the read depth of PA14 WT (Figure 1B and Figure S2B), the isolated clone (PA14*full) had far greater coverage across the prophage region (Figure 1C and Figure S2C) much like the evolved population (Figure S1), demonstrating that the *pf5r* mutation is responsible for the phage activation and that this trait can be horizontally transferred. Sequencing data also supported excision and circularization of the prophage genome, indicating prophage induction (Table S1), and qPCR of supernatants show that the increased read depth correlates with virion production (Figure S3). In summary, Pf5* prophage undergo increased replication and enable phage-encoded traits to be horizontally spread by superinfecting neighboring cells that have wildtype Pf5 prophage.

**Figure 1.**
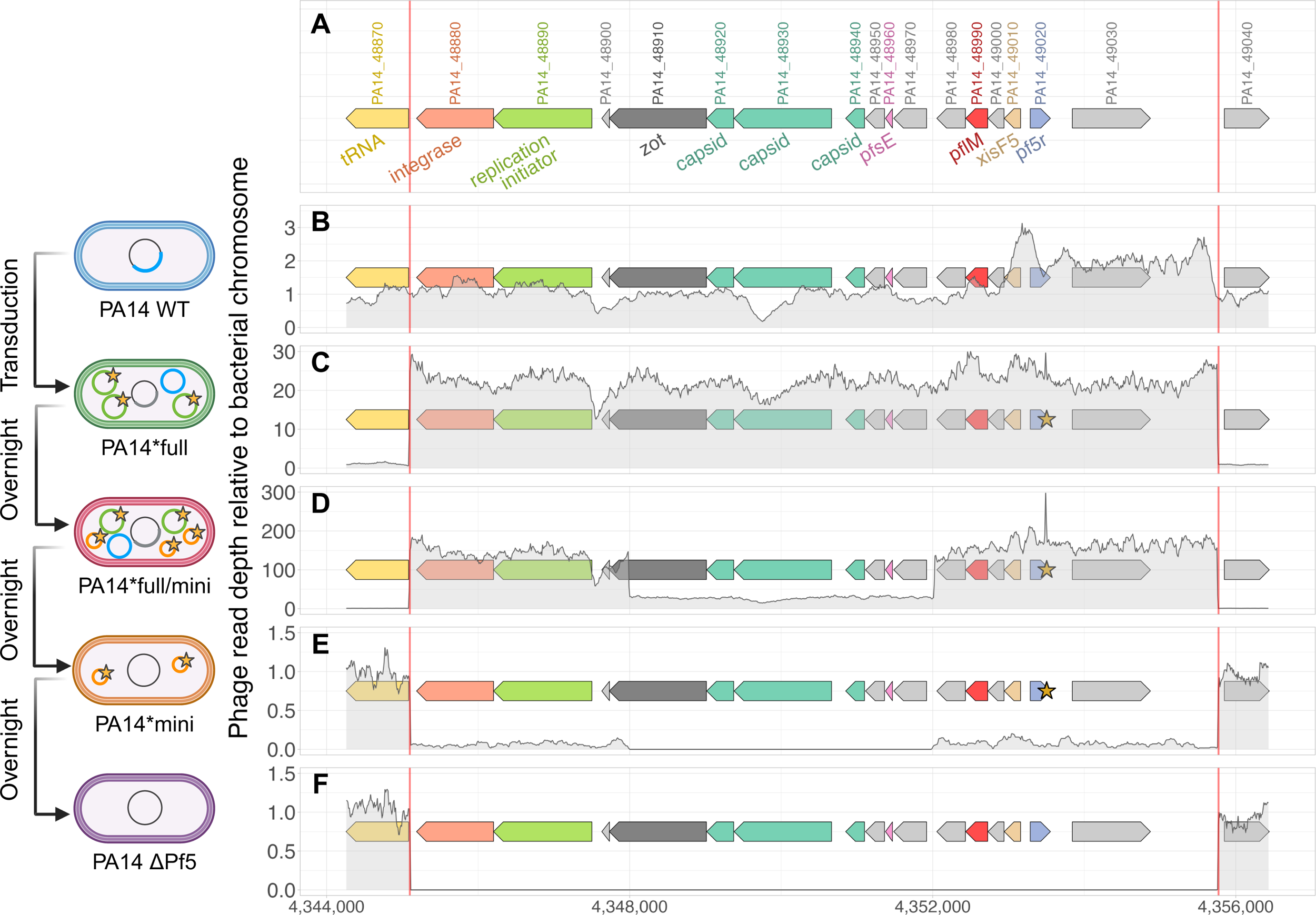
Hyper-replicative Pf5 phage (Pf5*full) increases intracellular phage diversity that enables invasion of incomplete miniphage. The y-axis denotes the read depth of the Pf5 prophage region relative to the average bacterial chromosome read depth 1000 bp upstream and 1000 bp downstream of the prophage region. The yellow star on the gene map denotes the position of the *pf5r* mutation. The red horizontal bars on the gene map denote the attachment sites (*attL* and *attR*) of the Pf5 phage. (A) Locus tags and annotations of the Pf5 prophage and flanking bacterial genes. Normalized read depths of (B) PA14 WT, (C) PA14*full, (D) PA14*full/mini, (E) PA14*mini, and (F) PA14ΔPf5 are shown. PA14*mini (in E) shows that some cells have already begun to lose Pf5 prophage completely as the read depth across the Pf5 prophage is lower than the read depth of the bacterial chromosome. (Created using R and BioRender.com.) See also Figure S1-4 and Table S1. Created in BioRender. Kubota, N. (2025) https://BioRender.com/kj3y0ut

### Cheater miniphages evolve within large Pf populations enabled by the defective Pf repressor

Defective Pf repressor genes have been associated with increased numbers of intracellular Pf phage, which increases the potential for the evolution of genetically diverse phage.^20^ Whole genome sequencing (WGS) of the newly isolated PA14*full showed increased copy numbers of both WT and mutated *pf5r* genes, indicating that both mutant and endogenous phage replication are upregulated (Table S1). Notably, in addition to the *pf5r* polymorphism, the genome lengths of Pf5 varied. Some PA14* isolates only carried full-length phages (i.e. PA14*full), others contained a mix of both full-length and smaller phage genomes (i.e. PA14*full/mini), and some bacterial clones only contained smaller phage genomes (PA14*mini). These smaller phage genomes, termed “miniphage,” have large deletions of multiple structural genes encoding capsid proteins and also the Pf superinfection exclusion (*pfsE*) gene (Figure 1D&E, Figure S2D&E, and Figure S4). We observed the emergence of miniphage multiple times among PA14* lineages (Figure 2) and even within the same bacterial clone (Figure 3), where the deletion lengths varied despite affecting the same structural genes. This high degree of parallelism indicates strong selection for these reduced phage genomes.

**Figure 2.**
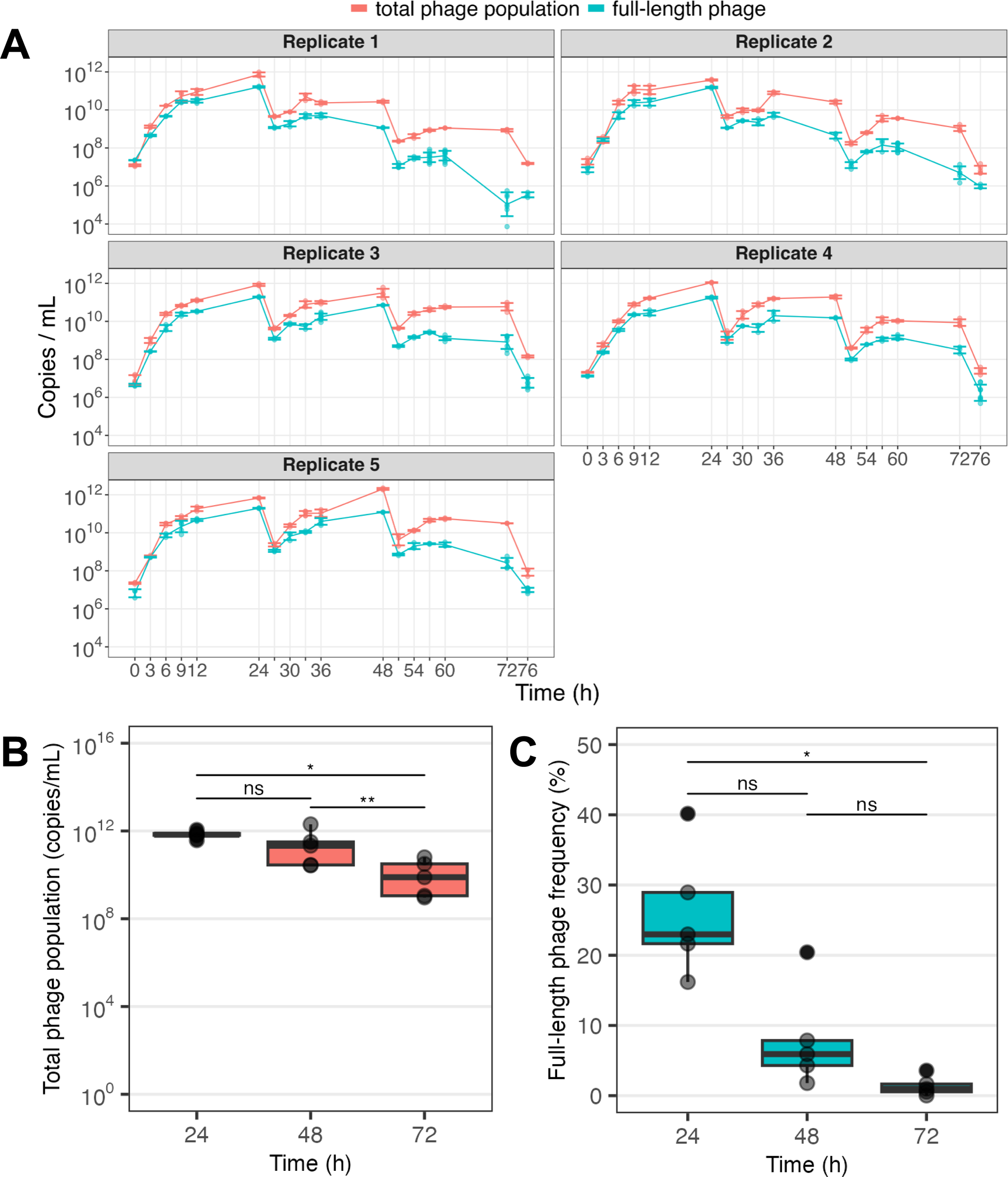
Miniphages emerge in PA14*full cultures within 24 hours and outcompete full-length phages while lowering total phage population. Five biological replicates of PA14*full cultures were grown in simplified CF media and tracked for 76 hours. Cultures were diluted 1:100 into fresh media every 24 hours. Total phage population was measured by qPCR of the *pflM* gene (in red), and the full-length phage population was measured by qPCR of phage structural genes (in blue). The miniphage population is inferred as the difference between these two measures. (A) Total phage population and full-length phage population over time. (B) Comparison of total phage population at 24, 48, and 72 hours. (C) Comparison of full-length phage frequency at 24, 48, and 72 hours. For (B) and (C), every point is the calculated average from each replicate and one-way repeated measures ANOVA with pairwise t-tests were performed with Bonferroni corrections to account for multiple comparisons (ns: p>0.05, *: p<0.05, **: p<0.01).

**Figure 3.**
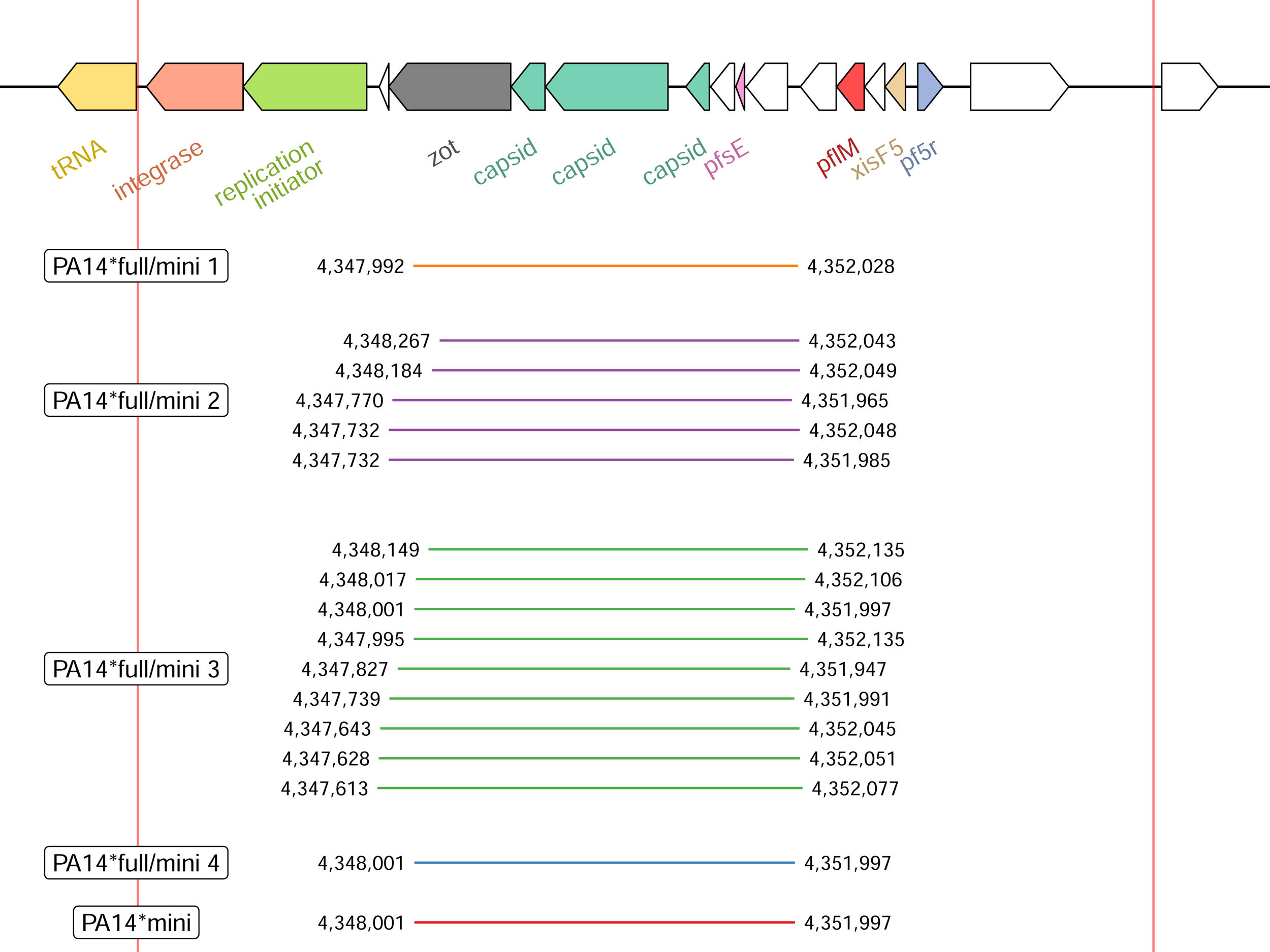
Genetically distinct miniphages emerge repeatedly following *pf5r* mutation. Five bacterial clones carrying miniphage(s) were isolated, and the red horizontal lines denote the deletion edges of the miniphage population within each clone. Some clones (i.e., PA14*full/mini 2&3) carry multiple miniphage genotypes, while others have only one (i.e., PA14*full/mini 1&4 and PA14*mini). Deletion edges vary in their exact position but are consistently located in the same region, with all five PA14*full/mini showing loss of the superinfection exclusion gene (*pfsE*) and structural genes such as capsid.

Miniphages can be considered cheaters because they do not produce structural phage proteins like capsid, which are public goods usable by any replicating phage. Under intense intracellular phage competition, phages with smaller genome sizes can potentially replicate faster than full-length phages and use such public resources to promote selfish replication. This exploitation can reduce phage populations in theory and was observed in independent studies of *de novo* miniphage invasion (Figure 2). Loss of the superinfection exclusion gene causes another conflict between full-length phages and miniphages, as superinfection by full-length phage benefits the miniphages but puts both full-length phages and the bacterial host at risk of additional phage invasion. Taken together, we observed that the emergence of the defective Pf repressor quickly leads to extreme levels of Pf reproduction, resulting in increased intracellular phage diversity, declining total phage numbers, and genome degradation due to intracellular phage competition.

### Pf5 repressor bacterial mutants release hyper-replicative phage that increase mutant fitness but lower population fitness

Filamentous phages uniquely do not require host cell lysis to reproduce and propagate.^8,9^ However, superinfective Pf4 phages have been linked to an autolysis phenotype in *P. aeruginosa* and can kill their bacterial host.^24,25^ To determine whether bacterial hosts containing different combinations of Pf5* genotypes were negatively affected by excess phage production, we compared their growth rates in isolation. The PA14* genotype grew significantly worse than WT, indicating that high phage production was costly (Figure 4A&B and Figure S5). Given the extreme cost of hyper-replicative Pf5, it seems paradoxical that defective *pf5r* would be favored during experimental evolution of PA14 populations maintained in large populations (>10^7^ cells) with efficient selection. We considered two non-mutually exclusive hypotheses: (1) the phage benefits from the defective repressor by increasing its replication rate, which enables horizontal spread in the bacterial population by superinfection, and/or (2) the bacterium benefits from *pf5r* mutant phage by being able to outcompete bacteria lacking them. We were able to horizontally transfer Pf5* phage into PA14 WT, so the former scenario is possible. However, PA14* also outcompeted the PA14 WT in direct competition and regardless of starting frequency (Figure 4C and Figure 5A), indicating that bacteria currently infected with Pf5* phage pay a lower cost for this infection than naïve cells. This demonstrates that PA14* bacteria somehow benefit from producing hyper-replicative Pf5 phage in direct competition with uninfected cells despite paying costs of reduced growth rates and yield.

**Figure 4.**
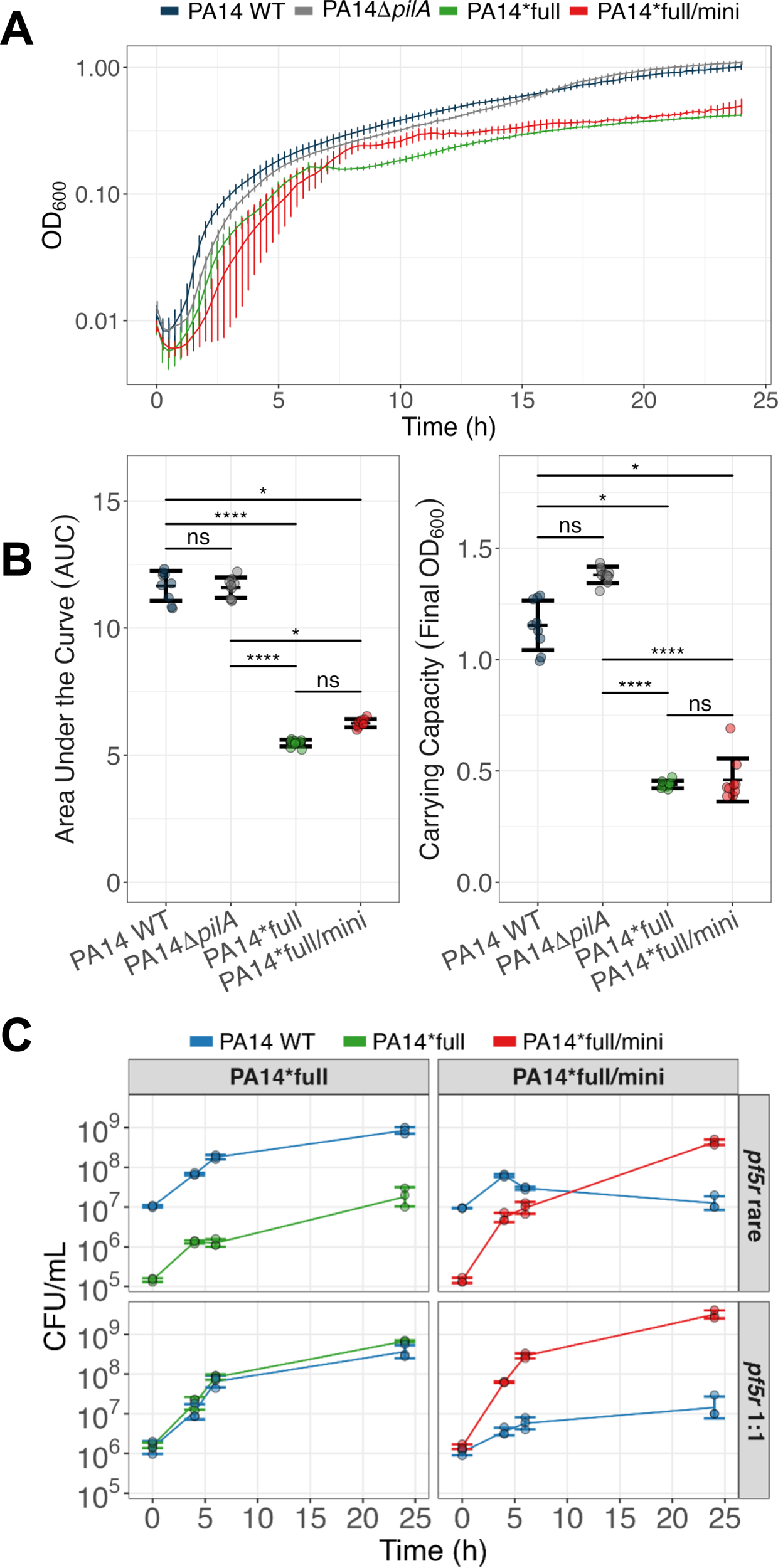
The *pf5r* mutation slows bacterial growth in monoculture but PA14*full/mini rapidly invades and outcompetes PA14 WT by suppressing WT growth. (A) Growth curve of all genotypes over 24 hours (OD_600_) represented as mean ± SD (*n* = 9 per strain); (B) Final OD_600_ and area under the curve (AUC) of the growth curves represented as mean ± SD (*n* = 9 per strain); (C) Top row: direct competitions when starting frequency of *pf5r* mutants was rare and PA14 WT common; bottom row: direct competitions when starting frequency of *pf5r* mutants and PA14 WT at 1:1 ratio, with outcomes measured as CFU/mL of each competitor over time. For each competition pairing, the averages of three biological replicates and their standard deviation are shown (Kruskal-Wallis test with Dunn’s where ns: p>0.05, *: p<0.05, ****: p<0.0001). See also Figure S5.

**Figure 5.**
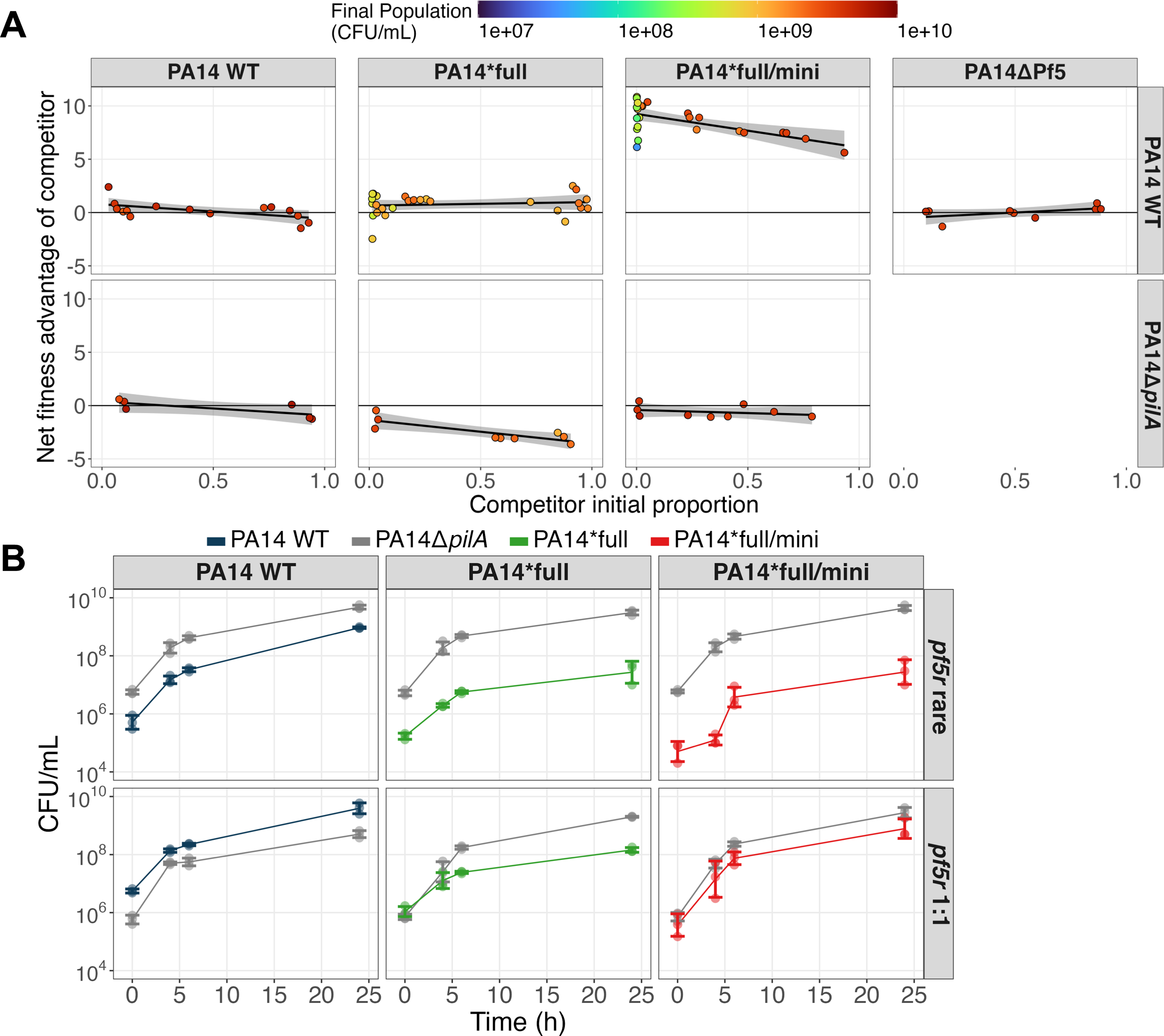
PA14*full and PA14*full/mini outcompete WT due to its greater susceptibility to superinfection. (A) Relative fitness (y-axis) of different bacterial strains (columns) containing mutant phage (with neutrality = 0) versus PA14 WT (top row) or PA14Δ*pilA* (bottom row) during competitions begun at a range of frequencies (x-axis). Every point denotes an independent competition, and the linear regression is the average net fitness across varying starting frequencies. PA14*full, and especially PA14*full/mini, outcompete WT at all frequencies but are outcompeted by PA14Δ*pilA* due to immunity of PA14Δ*pilA* to Pf5 superinfection. The loss of Pf5 (PA14ΔPf5) also eliminates any competitive advantage against PA14 WT. (B) Cell density (CFU/mL) of each competitor over time, in mixtures when *pf5r* mutants are rare (top row) relative to PA14 WT, and in mixtures where *pf5r* mutants and PA14 WT begin at 1:1 ratio (bottom row). For each competition pairing, the averages of three biological replicates and their standard deviation are shown.

We next tested whether the phages themselves, and not other physiological changes to PA14* induced by the phage, were responsible for suppressing PA14 WT growth. However, unlike Pf4 phage in PAO1, Pf5 phages do not form plaques on PA14 so we are unable to quantify phage via plaque assays. Instead, we competed the PA14*full and PA14*full/mini against PA14Δ*pilA*, which does not produce the phage receptor type IV pilus (TFP) and should be immune to Pf phage. PA14Δ*pilA* directly outcompeted both superinfected strains, confirming immunity (Figure 5A&B). Next, we tested whether phage-containing filtered supernatant from superinfected strains could inhibit WT growth as seen in direct competition. Supernatants of both PA14*full and PA14*full/mini suppressed WT growth but WT supernatant did not (Figure 6A). We repeated these supernatant experiments with PA14Δ*pilA* and found no growth defect (Figure 6A). These results further support the hypothesis that PA14*full and PA14*full/mini each suppress PA14 WT growth via Pf5* production. The fitness superiority of the TFP mutant that is immune to Pf5 superinfection shows that PA14* pays a cost for hyper-producing Pf5* phage, but the cost of Pf5* infection is greater to naïve PA14 WT than to PA14* (Figure 4C and Figure 5A).

**Figure 6.**
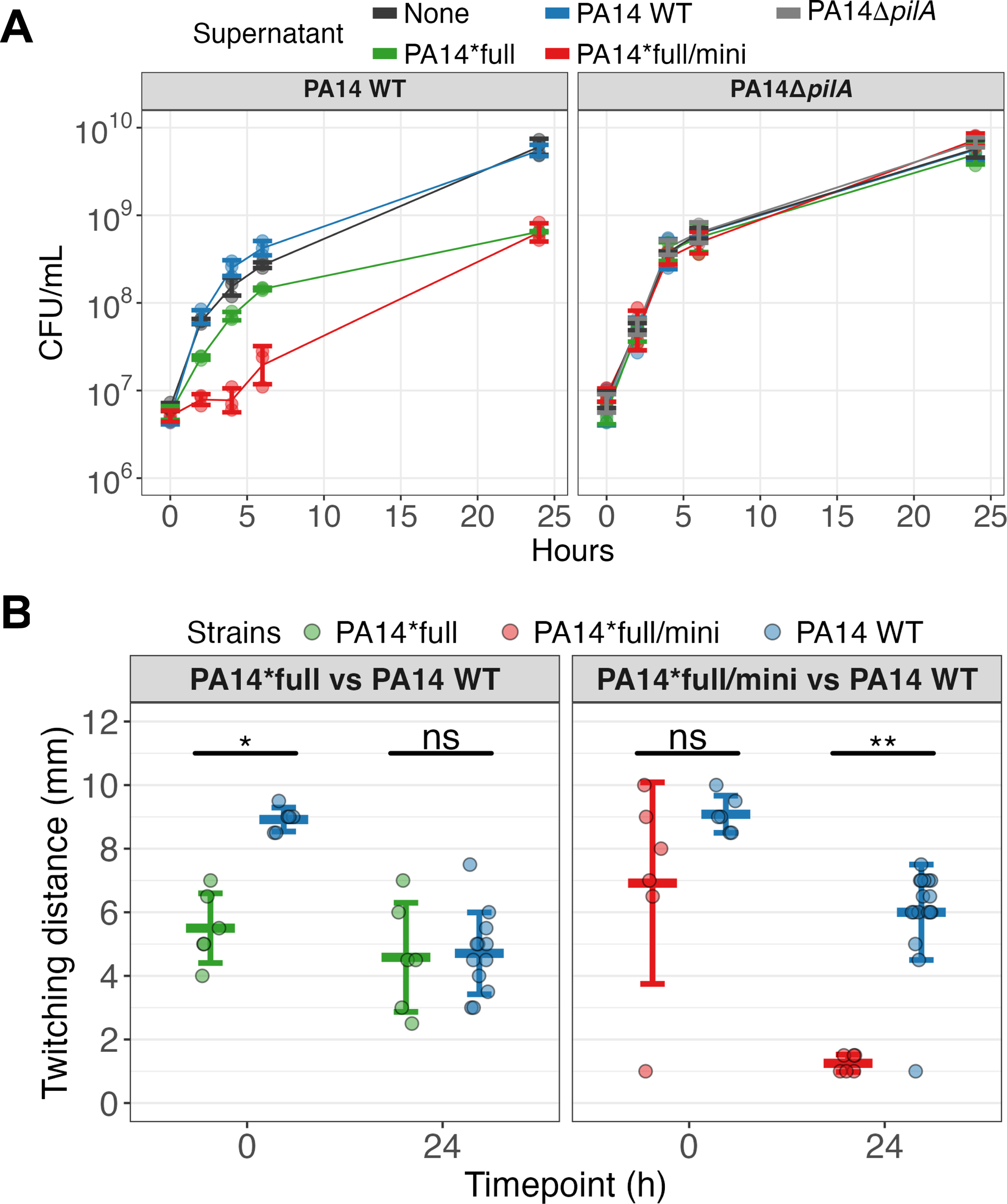
Pf5*full and Pf5*mini phages suppress PA14 WT growth. (A) Filtered supernatant from overnight cultures of PA14 WT, PA14Δ*pilA*, PA14*full, PA14*full/mini, or no supernatant was added to PA14 WT (left) or PA14Δ*pilA* (right) cultures. PA14*full/mini supernatant (containing both Pf5*full and Pf5*mini phages) suppresses PA14 WT growth but not PA14Δ*pilA*, which lacks the type 4 pili (T4P) phage receptor. Three different biological replicates were averaged, and the error bars show the standard deviation. (B) During competition against PA14 WT, PA14*full/mini exhibit reduced twitching motility mediated by TFP. Every point denotes the twitching distance of an individual colony that was isolated either at the beginning (0 h) or at the end (24 h) of the competition (n ≥ 6 for each strain in each competition; mean ± SD and Mann– Whitney U test with Bonferroni correction where ns: p>0.05, *: p<0.05, **: p<0.01.

### Bacteria with both hyper-replicative full-length phage and miniphage survive superinfection by downregulating the phage receptor

How PA14* seemingly tolerates Pf5* phage infection better than newly infected WT is unclear. Further, the PA14*full/mini genotype outcompetes WT better than PA14*full, despite both producing high levels of Pf5. We hypothesized that PA14*full/mini can better survive in the presence of Pf5* phage because this genotype can downregulate TFP function better than PA14*full, thereby becoming more phage-resistant. To measure TFP function, we assayed the extent of bacterial twitching motility, known to be mediated by TFP^26,27^, as a proxy. While PA14*full showed reduced twitching relative to PA14 WT prior to competition, PA14*full/mini became even less motile, indicating improved immunity to external Pf5* phage (Figure 6B). This ability to downregulate TFP to a greater extent provides an explanation for the greater fitness of the bacterium containing a mixture of full-length and miniphage relative to WT.

As noted previously, we observed multiple independent origins of miniphages in cells with defective Pf repressor genes (Figure 2 and Figure 3), but remarkably the deletion boundaries differed by only a few nucleotides (Figure 3). Each miniphage lost not only genes encoding structural proteins but also the *pfsE* gene encoding a superinfection exclusion protein (PA0721 in PAO1 and PA14_48960 in PA14).^7,21^ Many temperate phages encode superinfection exclusion mechanisms to protect their host from other phages and to decrease phage intracellular competition.^28,29^ PfsE prevents superinfection by binding to PilC in TFP, which prevents pilus extension and stops phages like Pf from binding to TFP.^7^ For miniphages lacking structural genes like capsid, losing PfsE allows their host to become superinfected with “cooperative” full-length Pf phage, thereby rescuing the miniphage from being lost by complementation. However, as suggested above, *pfsE* loss generates two new levels of conflict: first, between miniphage and full-length Pf phage by increasing phage intracellular competition via superinfection, and second, between miniphage and the bacterial host by increasing vulnerability of the bacterial host to other phages. Recurrent miniphage evolution clearly destabilizes the balance of fitness between Pf and their bacterial hosts.

### Bacteria with defective Pf repressor can rapidly lose Pf prophage due to intracellular phage competition

Rampant phage replication causes acute competition for host resources and public goods involved in replication. Consequently, cheater miniphages that do not produce capsid proteins experience increased fitness at the expense of cooperative, full-length phages. WGS of PA14* revealed that the intracellular phage population in a single bacterial colony can be remarkably diverse, with some cells carrying different mixtures of Pf5 WT, Pf5*full, and Pf5*mini phages including multiple genotypes within each phage class (Table S1). We grew five independent PA14*full cultures and used qPCR of *pfsE* as a proxy for total phage quantification and the region spanning the *zot* gene and capsid genes as a proxy for full-length phage quantification. The miniphage population, inferred as the difference between the two prior quantities, rose extremely rapidly within clonal bacterial populations, becoming a significant portion of the phage population within 24 hours or approximately <7 population-wide bacterial generations (Figure 2). This rapid miniphage invasion has also been reported for Pf4 infecting PAO1.^30^ In each case, the emergence of miniphage led to the decline of the total phage population over time, with full-length phages declining more rapidly than miniphages (Figure 3). Eventually, through subsequent uneven vertical inheritance of phages, we observed that some bacteria only inherit miniphages (Figure 1E and Table S1). Once this happens, miniphages can no longer leave the cell, and in following cell divisions, the bacteria lose the phage completely (PA14ΔPf5; Figure 1F). This process defines a Tragedy of the Commons for the Pf phages as competition leads to the complete loss of full-length phage genomes that can produce capsid and thus the phage population goes locally extinct. Nonetheless, WGS of PA14ΔPf5 shows that the *pf5* attachment sites in PA14ΔPf5 remain intact, indicating the possibility for reinfection. We competed PA14ΔPf5 directly against PA14 WT and found it lost the competitive advantage of the original PA14* genotype, implying that the phage-free state is ecologically unstable and prone to reinfection and re-initiation of this life cycle (Figure 5A).

## Discussion

Molecular parasites such as phages can increase their evolutionary potential and fitness by expanding their population size. In such conditions, multiple genotypes can arise and cooccur within a host cell, including defective variants that can be rescued and persist through complementation by complete phage genomes.^31,32^ However, large parasite populations increase the burden on the host, leading to greater conflict between host and parasite and also greater intracellular conflict between parasites as competition for host resources intensifies. In addition, genetic and phenotypic diversification of these molecular parasites can affect host fitness and introduce diversity between infected cells. Many bacteria carry prophages in their genomes, and *P. aeruginosa* and their widespread Pf prophage are no exception.^10^ While some aspects of bacteria and phage interaction can be mutualistic, such as the provision of antibiotic resistance genes and immunity against other parasites, this pact can quickly shift towards parasitism. Here, we show that bacteria can be at the mercy of rapid changes in their prophage population developing in mere hours that slow bacterial reproduction but transiently increase relative bacterial fitness. Specifically, we report that Pf phages that evolved a defective Pf repressor provide the bacterial host with the ability to use Pf as a weapon against other bacterial competitors, but that this state is unstable: the emergence and spread of cheater miniphages result in a phage Tragedy of the Commons leading to subsequent phage extinction.

Viruses can cheat other viruses infecting the same host by not contributing to public goods production.^33^ For example, viruses lacking capsid genes do not contribute to capsid production but can use capsid proteins produced by co-infecting “cooperative” viruses, thereby selfishly propagating at the expense of the cooperators. Filamentous phages, including M13 and f1 that infect *E. coli* and Pf4 in PAO1, have been observed to lose large parts of their genome creating cheater miniphages.^30,34–37^ These miniphages have been reported in *P. aeruginosa* Pf previously^19,30^ and one study created miniphages by deleting structural genes^38^, but otherwise, Pf miniphages have not been actively studied. When phage replication is rampant, miniphages arising initially at low frequency gain a large fitness advantage as they have many cooperators to exploit. In Pf4 phages of *P. aeruginosa*, deletion of the Pf4 repressor (*pf4r*) gene was observed to increase phage production^17^ and to produce Pf superinfective variants that increase intracellular phage diversity.^18,20^ These superinfective Pf variants were reported to kill their host cell and play roles in bacterial biofilm dispersal and in phenotypes such as small colony variants and bacterial virulence.^18,22,24,25,39,40^ In this study, we show that the emergence of superinfective Pf5 variants in strain PA14 leads to the subsequent invasion of Pf miniphages and a rapid sequence of events affecting bacterial fitness. We were able to detect these dynamics by sequencing whole genomes of bacterial clones and populations over short intervals and focusing on spikes in read depth as well as reads that support circularization of the Pf5 prophage genome. This study presents both a conceptual model and analytical methods for interpreting the genomes of bacteria carrying filamentous phage, which may reveal ongoing interactions in their intracellular virome.

In summary, we demonstrate that intracellular phage competition can favor rapid expansion of the phage population both within and among neighboring host bacteria, but this evolution is short-sighted, as short-term gains in phage fitness lead to their eventual local extinction. From the perspective of the bacterial cell, its fitness is aligned with the hyper-replicative phage at the outset, because the cost of phage replication is outweighed by increased relative fitness against other cells becoming superinfected. The emergence of cheater phages also benefits the host by improving immunity against further superinfection, but it dooms the phage to a Tragedy of the Commons and extinction. Phage loss eliminates any burdens of phage replication for the host but it also eliminates the relative fitness advantage of infected cells, leaving the bacterial population susceptible to invasion.

Superinfective variants of Pf can readily appear and have been documented to be an important part of *P. aeruginosa* lifestyle relevant for a broad array of clinically important bacterial phenotypes, especially in biofilm.^14,19,20,24,41^ This study shows that the emergence of such hyper-replicative and superinfective variants favors the invasion of miniphages both within and among host cells. But since Pf phage population structure can shift in a manner of hours and affect bacterial fitness, antimicrobial resistance, defense against host immunity, and aspects of pathogenicity^10,11,13,15^, our study reveals a potentially common yet overlooked driver of *P. aeruginosa* evolution and potentially of all bacteria harboring filamentous phages.^42^

## Supporting information

KeyResources

## RESOURCE AVAILABILITY

### Lead contact

Requests for further information and resources should be directed to and will be fulfilled by the lead contact, Vaughn S. Cooper.

### Materials availability

All strains present in this study are available upon request.

### Data and code availability

- All raw reads from the whole genome sequences have been deposited with links to BioProject accession number PRJNA692838 and PRJNA1226961 in the NCBI BioProject database (https://www.ncbi.nlm.nih.gov/bioproject/).
- All data and code used to analyze variant calls, read depth, growth curves, competition assays, and twitch assays can be found at https://github.com/NanamiKubota/pf_cheater_phage.
- Any additional information required to reanalyze the data reported in this paper is available from the lead contact upon request.

## Supplemental information

Document S1. Figures S1-S5 and Table S1-S3.

## Acknowledgments

We thank members of the Cooper lab, the Pat Secor lab, and thesis committee members (Drs. Daria Van Tyne, Will DePas, Graham Hatfull, and Pat Secor) for valuable feedback and support. We also thank the reviewers for their helpful comments in improving this manuscript. Research was supported in part by the Ruth L. Kirschstein Predoctoral Individual National Research Service Award (1F31AI179118-01 to NK), by NIH U19AI158076 to VSC, and by the Pennsylvania Department of Health PA CURES Grant #4100085725. The content is solely the responsibility of the authors and does not necessarily represent the official views of the National Institutes of Health or the Pennsylvania Department of Health.

## Author contributions

Conceptualization, writing, review, and final approval of the manuscript were done by NK, MRS, and VSC. Experiment and bioinformatics analyses were done by NK and MRS.

## STAR Methods

### EXPERIMENTAL MODEL

*Pseudomonas aeruginosa* UCBPP-PA14 *pf5r* mutant clones were originally isolated from an evolution experiment, but these mutants contained secondary mutations.^16,23^ To create a *pf5r* mutant in the PA14 WT background (PA14*full), we co-cultured an unmarked, evolved PA14 *pf5r* mutant with *lacZ*-marked PA14 WT overnight and screened blue colonies for mobilized Pf5 *pf5r* phage via qPCR across the attachment sites (attB and attP) for excised and circularized Pf5 phage, respectively, with *gyrB* as the housekeeping gene (see Table S3 for list of primers). Colonies with high copy number for attB and attP were analyzed by whole genome sequencing (WGS) to confirm presence of the *pf5r* mutation via horizontal gene transfer.

PA14*full/mini and PA14*mini were isolated by streaking PA14*full on 1/2 Tryptic Soy (Tsoy) Agar and screening colonies for capsid deletion via PCR. PA14ΔPf5 was isolated by streaking PA14*mini on 1/2 Tsoy agar and screening across two conserved phage genes in Pf5*mini (i.e., PA14_48980-PA14_489900). All mutants were verified via WGS. PA14Δ*pilA* mutant was generated through two-step allelic exchange^47^ and confirmed via WGS. Detailed nomenclature of bacteria and phage strains can be found in Table S2. Bacteria were cultured in glass tubes at 37°C on a roller drum with M9 minimal media supplemented with nutrients found in cystic fibrosis lungs (i.e. CF media) as previously described.^16^ In short, the media comprised of an M9 salt base (0.1 mM CaCl_2_,1.0 mM MgSO_4_, 42.2 mM Na_2_HPO_4_, 22 mM KH_2_PO_4_, 21.7 mM NaCl, 18.7 mM NH_4_Cl), 11.1 mM glucose, 10 mM DL-lactate (Sigma-Aldrich L1375, CAS 72-17-3), 20 mL/liter MEM essential amino acids (ThermoFisher 11130051), 10 mL/liter MEM non-essential amino acids (ThermoFisher 11140050), and 1 mL/liter each of Trace Elements A, B, and C (Corning MT99182CI, MT99175CI, MT99176CI).

## METHOD DETAILS

### Whole genome sequencing and analysis

DNA of all bacteria strains were extracted using the Qiagen DNeasy Blood & Tissue Kit and WGS using Illumina NextSeq550 with a resolution of 200 Mbp or between 3-14 million reads per sample. Reads were checked in fastQC^43^ for quality control. The PA14 WT reads were mapped against the PA14 from the Pseudomonas Genome Database^48,49^ (https://www.pseudomonas.com/strain/show/109) using the variant caller breseq v0.39.0^44^ with the consensus mode default parameters. Then the gdtools program within breseq was used to create a new PA14 reference genome that contains mutations present in our PA14 lab wildtype strain. Raw reads of all other samples were mapped against this new reference. Read depth was calculated by averaging the reads mapped to a 10 bp window across the bacterial genome using bedtools^45^ (v2.26.0) and samtools^46^ (v1.21). The raw reads have been deposited with links to BioProject accession number PRJNA692838 and PRJNA1226961 in the NCBI BioProject database (https://www.ncbi.nlm.nih.gov/bioproject/). Read depth plots were created using R^50^ (v4.4.0), RStudio^51^ (v2024.04.2+764), tidyverse^52^ (v2.0.0), ggplot2^53^ (v3.5.1), and gggenes^54^ (v0.5.1).

### Growth curves

Growth curves were conducted in 96-well plates with each well seeded with 198µL of CF media and 2µL overnight bacterial culture. Optical density was read at 600nm (OD_600_) every 15 minutes over the course of 24 hours at 37°C. OD_600_ was graphed in R, and the growth rate, area under the curve (AUC), and the final OD_600_ was calculated using the Growthcurver package.^55^

### Competition assays

Overnight cultures of each competitor strain were grown in individual tubes in CF media at 37°C prior to the competition. Competitors were paired such that one competitor was *lacZ*-marked while the other was unmarked. Approximately 1mL of overnight cultures were pelleted and washed with 1xPBS to remove phages that may be present in the supernatant. The paired competitors were mixed at varying ratios so that the total volume of the co-culture was 50 µL and this was seeded into 5 mL of fresh CF media to make a 1:100 dilution. For competitions where CFU/mL were measured multiple times between timepoints 0 h and 24 h, a 1:1 volume ratio was used to seed competitions where *pf5r* mutants were rare relative to their competitors (either PA14 WT or PA14Δ*pilA*). For competitions where *pf5r* mutants were approximately 1:1 in starting CFU/mL with competitors, a 1:9 volume ratio of competitor-to-*pf5r* culture was used.

The co-cultures were then vortexed and starting CFU/mL (t = 0 h) was quantified by serial diluting the co-cultures in 1xPBS and spread plating the dilutions onto 1/2 Tryptic Soy Agar plates supplemented with X-gal. The co-cultures were then incubated at 37°C on a roller drum and CFU/mL was measured at various timepoints using plate dilutions up until 24 h post-inoculation. Blue and white colonies were counted, and selection rate constant (*r*) was calculated by taking the difference in Malthusian parameter of the two competitors^56^: *r* = 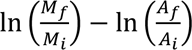, where *M_i_* and *A_i_* is the mutant or ancestor CFU/mL at 0 h post-inoculation, respectively, and *M_f_* and *A_f_* is their CFU/mL at 24 h post-inoculation, respectively.

### Pf5* phage treatment on PA14 WT and PA14Δ*pilA*

Overnight cultures of PA14 WT, PA14Δ*pilA*, PA14*full, and PA14*full/mini were grown in CF media at 37°C. Approximately 1 mL of PA14*full and PA14*full/mini overnight cultures were passed through a 0.20 µm filter into separate microcentrifuge tubes. Then into 5 mL of fresh CF media, 50 µL of the filtrate and 50 µL of the PA14 WT or PA14Δ*pilA* overnight were added and vortexed. The CFU/mL of the initial culture (0 h) was measured by taking 20 µL of the seeded culture and doing ten-fold dilutions in a 96-well plate with 1xPBS. 100 µL of the diluted culture was plated onto 1/2 Tsoy supplemented with X-gal before incubating the plates overnight. The culture tubes were removed from the incubator at 2 h, 4 h, 6 h, and 24 h to measure CFU/mL as described earlier.

### Twitching assay

Twitching motility were measured following an established protocol.^27^ Briefly, a single colony from the competition experiments (either the 0 h timepoint or 24 h timepoints) were picked using a pipette tip. The colony color was used to distinguish PA14*full and PA14*full/mini (blue; *lacZ*-marked) from PA14 WT (white; unmarked). The center of the 1% LB-Lennox Agar was stabbed using the pipette tip, making sure that the tip touched the bottom of the petri dish. Then, the plates were incubated at 37°C overnight in a plastic container with a lid and lined with damp paper towels to prevent the plates from drying. The following day, plates were flooded with cold TM developer solution for 30 minutes to make the bacteria opaque. Twitching distance was measured from the center of the plate where the stabs were made to the edge of the front using a ruler.

### Pf5 virion quantification via qPCR

Five independent overnight cultures of PA14*full were grown in 5 mL of CF media at 37°C. From each overnight cultures, 1 mL of the overnight cultures were pelleted in separate microcentrifuge tubes and the supernatant was discarded. The pellet was washed with 1xPBS to remove any residual phages and pelleted again to discard the supernatant before being resuspended in 1 mL 1xPBS. Then 65 µL of the resuspended culture was added into 6.5 mL fresh CF media to make 1:100 dilution culture. 500 µL of the new culture was collected passed through a 0.20 µm filter into a new microcentrifuge tube. The remaining culture was incubated at 37°C. Supernatants were collected multiple times in a similar manner over the course of 76 hours. Every 24 hours, 65 µL of culture was passaged to 6.5 mL fresh CF media. The filtered supernatant underwent DNase I treatment following previously published protocols with minor modifications to align with manufacturer’s instructions.^7,19^ Briefly, 89 µL of the filtered supernatant was added to a new microcentrifuge tube with 1 µL DNase I (NEB M0303S) and 10 µL DNase I Reaction Buffer (NEB B0303S) to create a 100 µL total reaction volume. The supernatant with the DNase I was incubated at 37°C for 1 hour, followed by a DNase inactivation step at 75°C for 10 minutes. 2 µL of the treated supernatant was qPCR using primers that amplified the *pflM* gene, zot/capsid gene, and *gyrB* (negative control; see Table S3 for list of primers). No *gyrB* amplification was detected, indicating that DNA not protected by the phage capsid was digested by the DNase I. For the standard curves, PCR amplicons produced by the qPCR primers with PA14 WT DNA were used. The PCR amplicons underwent gel extraction to remove any impurities before standard curves were created. Standard curves were used to quantify total phage population (via *pflM* amplification) and full-length phage numbers (via *zot*-capsid amplification). Miniphage numbers are inferred from the difference between full-length phage and total phage population. Supernatants from overnight cultures of various PA14 strains were also treated using the same filtration, DNase treatment, and qPCR protocol.

## QUANTIFICATION AND STATISTICAL ANALYSIS

All statistical analyses were conducted in R^50^ (v4.4.0) and RStudio^51^ (v2024.04.2+764) unless otherwise noted. Read depth calculations were done using bam output files provided by breseq^44^ (v.0.39.0) and measured at 10 bp windows using bedtools^45^ (v2.26.0) and samtools^46^ (v1.21). Detailed information about the statistical test is provided in the figure legend. Data is presented as mean ± SD unless otherwise noted. All experiments were conducted with at least three biological replicates unless otherwise noted.

