## SupplementalFigs-Tables for "Filamentous cheater phages drive bacterial and phage populations to lower fitness"

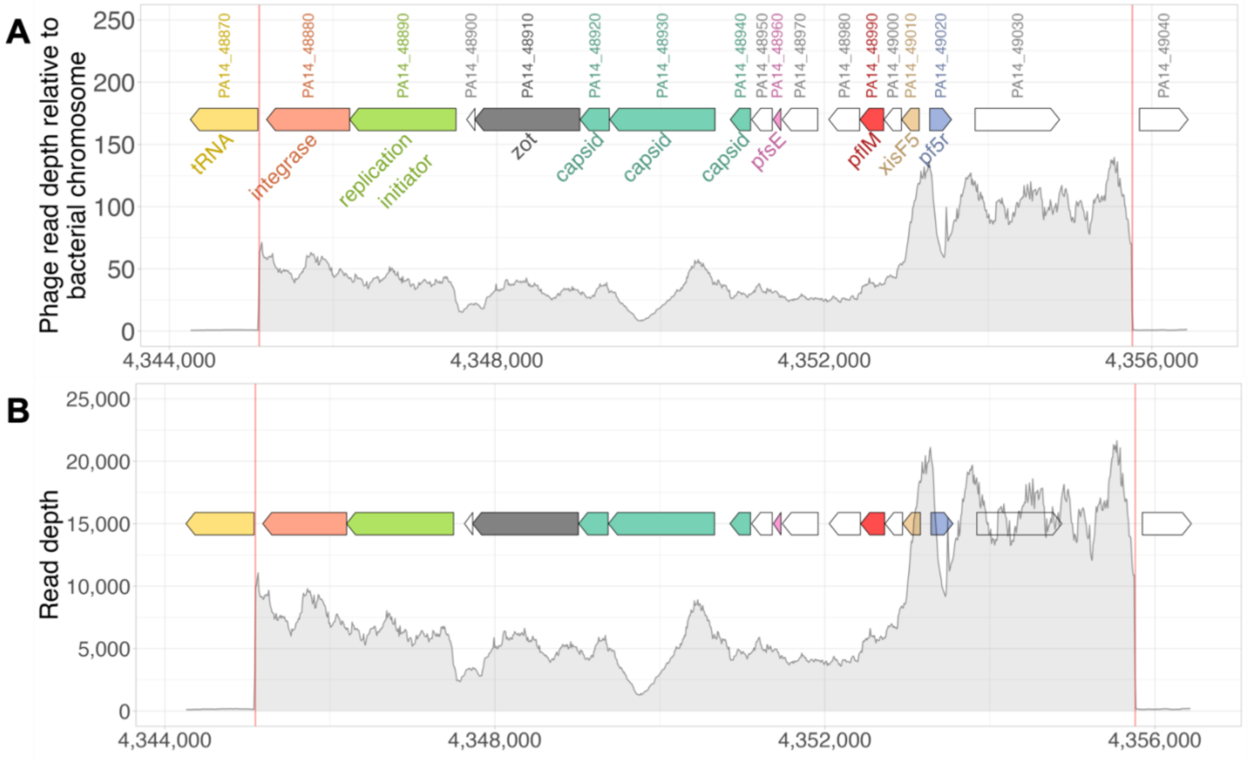


**Figure S1. Read depths of the prophage region are elevated in the evolved population containing Pf5 pf5r mutant phage. Related to Figure 1.** Read depth was measured in 10 bp increments. (A) The read depth across the Pf5 prophage region was normalized to the read depth of the bacterial regions flanking 1 kb upstream and downstream the prophage. (B) Raw read depth values without any normalization.

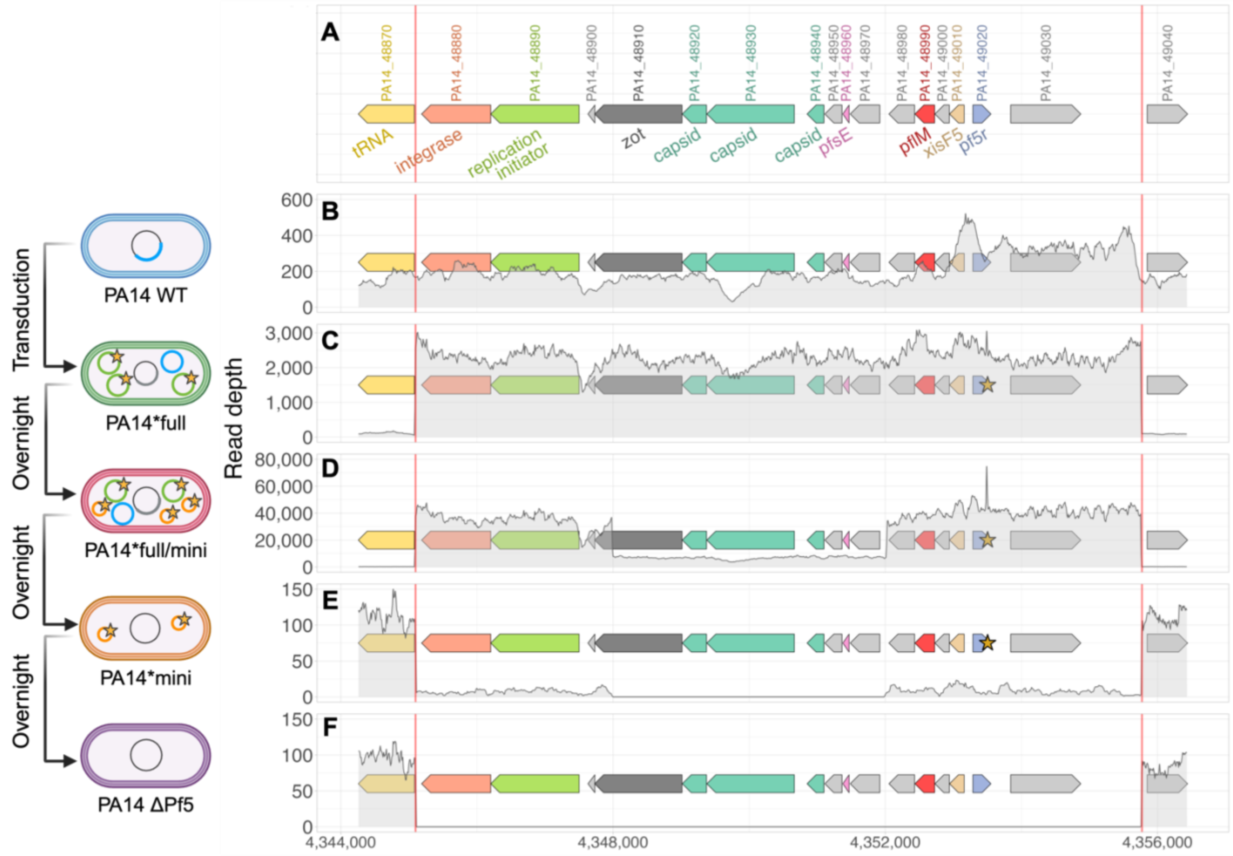


**Figure S2. Raw read depth of Pf5 prophage region of cells with different viral populations relative to the flanking bacterial chromosome. Related to Figure 1.** This is the same data from Figure 1, but the reads are not normalized. Created in BioRender. Kubota, N. (2025) https://BioRender.com/qjdalg2

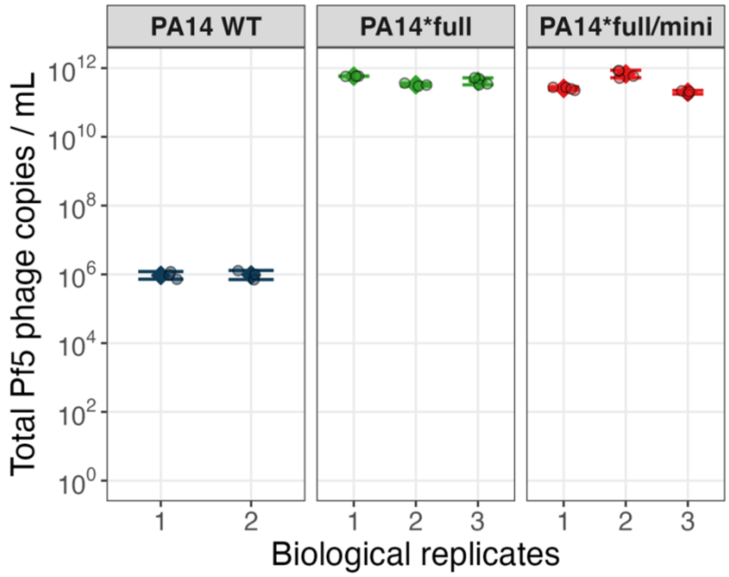


**Figure S3. Prophage repressor mutant produces more Pf5 phage than PA14 WT. Related to Figure 1.** Supernatant of independent overnight cultures (biological replicates) were filtered, DNase treated, and analyzed by qPCR of *pflM* for total Pf5 phage population. Each biological replicate has at least 3 technical replicates (circles). Data are shown as mean ± SD. PA14*mini and PA14ΔPf5 are not shown as Pf5 phages were not detected in the supernatant.

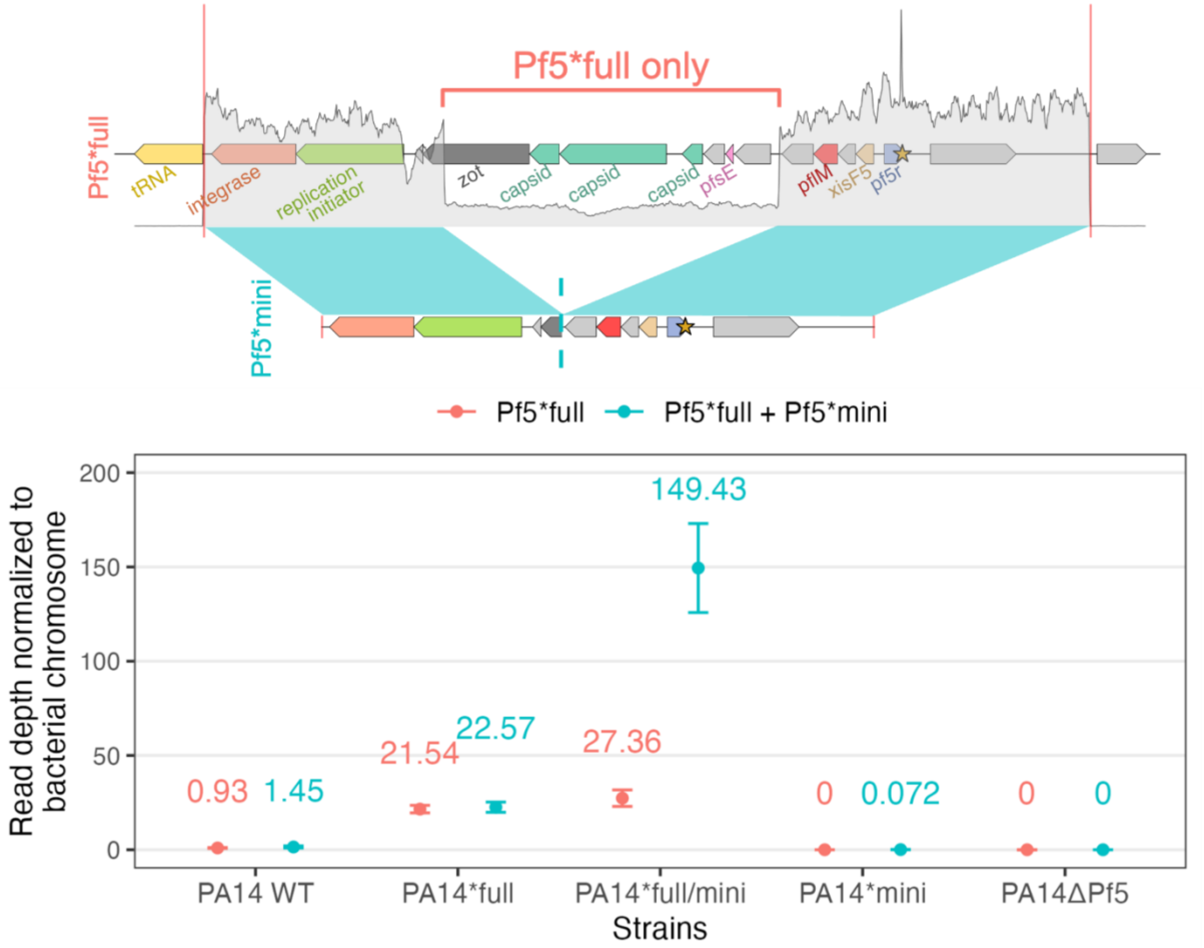


**Figure S4. Read depths of the phage genome normalized to the bacterial chromosome show that the phage copy number increases when *pf5r* is mutated and increases again when both Pf5*full and Pf5*mini are present. Related to Figure 1.** Read depth was measured in 10 bp increments and then normalized to coverage of the surrounding bacterial chromosome before calculating the mean coverage and standard deviation for each Pf5 region. Created in BioRender. Kubota, N. (2025) https://BioRender.com/f9wmf1y

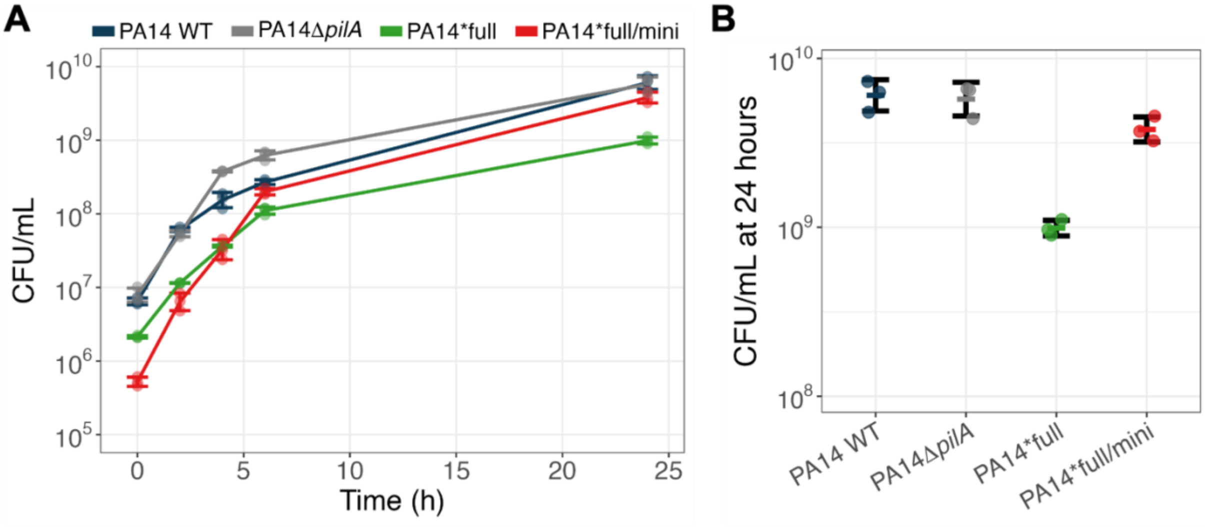


**Figure S5. CFU/mL measurements of the bacteria correlate with $OD_{600}$ readings. Related to Figure 3.** (A) CFU/mL of each strain over 24h of growth and (B) CFU/mL comparison at 24 hours (mean ± SD, *n* = 3).

| **Bacterial strain** | **mutated *pf5r*** | **WT *pf5r*** | **Pf5 excision** | **Pf5 circularization** | **miniphage** |
|---|---|---|---|---|---|
| PA14 WT | 0% | 100% | 0% | 2.7% | 0% |
| PA14*full | 38.1% | 61.9% | 16.7% | 97.2% | 0% |
| PA14*full/mini | 80.6% | 19.4% | 5.8% | 99.7% | 79.4% |
| PA14*mini | 94.1% | 5.9% | 97.1% | 0% | 100% |
| PA14ΔPf5 | 0% | 0% | 100% | 0% | 0% |

**Table S1. Frequency of reads supporting *pf5r* mutation shows that clonal bacteria carry both mutated and wild-type (WT) copies of the *pf5r* gene until Pf5*mini phages drive the phage population to a Tragedy of the Commons. Related to Figure 1.** WT *pf5r* frequency is inferred by subtracting the frequency of mutated *pf5r* reads from 100%, except for PA14ΔPf5 where Pf5 phage is extinct.

| Organism | Strain | Description | Accession | Origin |
|---|---|---|---|---|
| Bacteria | PA14 WT | *Pseudomonas aeruginosa* UCBPP-PA14 (also known as PA14) wildtype strain with wildtype Pf5 prophage and an intact *pf5r* gene. | | Scribner et al.[S1] |
| | PA14*full | PA14 wildtype infected with full-length Pf5 phage with defective *pf5r* gene (i.e. Pf5*full phage). | SAMN23928097 | Scribner et al.[S1] |
| | PA14*full/mini | PA14 infected by both Pf5*full and Pf5*mini phage. | SAMN46967948 | This study |
| | PA14*mini | PA14 only infected by defective *pf5r* miniphage (Pf5*mini). | SAMN46967946 | This study |
| | PA14ΔPf5 | PA14 that lacks Pf5 phages/prophages but with an intact attB site. | SAMN46967947 | This study |
| Phage | Pf5 WT | Wildtype Pf5 phage, usually found integrated into PA14 WT chromosome as a prophage and rarely induced. | | Scribner et al.[S1] |
| | Pf5*full | Full-length Pf5 phage with defective *pf5r* gene isolated from the final evolved population of planktonic no drug cultures grown in CF media from a previous evolution experiment. | | Scribner et al.[S1] |
| | Pf5*mini | Pf5 miniphage with defective *pf5r* gene that lost several genes, including the Pf5 capsid genes. | | This study |

**Table S2. Detailed nomenclature of bacteria and phage strains in this study. Related to STAR Methods.**

| Name | Sequence (5’-3’) | Purpose | Reference |
|---|---|---|---|
| PA14*gyrB*-f | cacagcatccagcgatacaag | qPCR bacteria housekeeping gene | Li et al. [S2] |
| PA14*gyrB*-r | cgcgttgctttcgatgaagt | qPCR bacteria housekeeping gene | Li et al. [S2] |
| zot-capsid-qPCR-F | gcctgaggtcttgtaggagc | qPCR full-length phage | This study |
| zot-capsid-qPCR-R | gccgttcattgggaggtgat | qPCR full-length phage | This study |
| pflM-qPCR-F2 | gccatcttccagatagggca | qPCR total phage population | This study |
| pflM-qPCR-R2 | cagtccctactacctgcacc | qPCR total phage population | This study |
| pf5 capsid F 1 | cgcacgcttttcaatcctca | PCR Pf5 capsid gene | This study |
| pf5 capsid R 1 | ctatctctcgctgttcgcgg | PCR Pf5 capsid gene | This study |
| pf5 capsid del F 1 | attcggaatcggggacgatg | PCR Pf5 capsid gene loss | This study |
| pf5 capsid del R 1 | gacgacgttgcgatagggat | PCR Pf5 capsid gene loss | This study |
| Pf miniphage F 1 | ggtaacgatccctatcgcaa | PCR Pf5 miniphage | This study |
| Pf miniphage R 1 | aagtactggcacgttgtgta | PCR Pf5 miniphage | This study |
| PA14-pilA-Up-R | aatatgcctgccctgactgc | *pilA* deletion cloning | This study |
| PA14-pilA-Up-F | gccagtgccaagcttgcatgcctgcaggtcgactctagaggcctcgcctgatagacaaca | *pilA* deletion cloning | This study |
| PA14-pilA-Down-R | agctatgaccatgattacgaattcgagctcggtacccgggtcgccacaacggaactactc | *pilA* deletion cloning | This study |
| PA14-pilA-Down-F | gcagtcagggcaggcatattgcatcttgcatgccaacctg | *pilA* deletion cloning | This study |
| M13 fwd | gtaaaacgacggccagt | *pilA* deletion cloning | universal primer |
| M13 rev | caggaaacagctatgac | *pilA* deletion cloning | universal primer |
| PA14-pilA-del-seq-F | ggccagcctgcagataagac | *pilA* deletion cloning | This study |
| PA14-pilA-del-seq-R | tcgatgatgatgccgagctt | *pilA* deletion cloning | This study |

**Table S3. List of primers used in this study. Related to STAR Methods.**
