## Supplementary material for "Filamentous cheater phages drive bacterial and phage populations to lower fitness": KeyResources

**Key resources table**

| REAGENT or RESOURCE | SOURCE | IDENTIFIER |
| --- | --- | --- |
| Bacterial and virus strains | | |
| *P. aeruginosa* PA14, WT | Scribner et al.^16^ | See Table S1 |
| *P. aeruginosa* PA14, WT *lacZ* | Lab stock | N/A |
| *P. aeruginosa* PA14, PA14*full | This study | See Table S1 |
| *P. aeruginosa* PA14, PA14*full/mini | This study | See Table S1 |
| *P. aeruginosa* PA14, PA14*mini | This study | See Table S1 |
| *P. aeruginosa* PA14, PA14∆Pf5 | This study | See Table S1 |
| Various Pf5 phage strains | This study | See Table S1 |
| *P. aeruginosa* PA14, PA14∆*pilA* | This study | N/A |
| *E. coli* S17-1 (λpir) | Laboratory of Catherine Armbruster | N/A |
| *E. coli* DH5a | Lab stock | N/A |
| Deposited data | | |
| Raw reads from whole genome sequencing of various *P. aeruginosa* infected with Pf5 phage | This study | BioProject PRJNA1226961 |
| Raw reads from whole genome sequencing of previous study’s *P. aeruginosa* evolution experiment | Scribner et al.^16^ / NCBI | BioProject PRJNA692838 |
| Processed data | This study | https://github.com/NanamiKubota/pf_cheater_phage |
| *P. aeruginosa* UCBPP-PA14 | NCBI | BioSample Accession SAMN02603591 |
| Oligonucleotides | | |
| Primers for cloning and PCR/qPCR, see Table S3 | This study | N/A |
| Recombinant DNA | | |
| pEX18Gm | NovoPro | Cat No. V006439 |
| Software and algorithms | | |
| fastQC v0.11.5 | Andrews^43^ | https://github.com/s-andrews/FastQC |
| breseq v0.39.0 | Deatherage and Barrick^44^ | https://github.com/barricklab/breseq |
| bedtools v2.26.0 | Quinlan and Hall^45^ | https://github.com/arq5x/bedtools2 |
| samtools v1.21 | Li et at.^46^ | https://github.com/samtools/samtools |
